# Longitudinal single-cell data informs deterministic modelling of inflammatory bowel disease

**DOI:** 10.1101/2023.10.27.561846

**Authors:** Christoph Kilian, Hanna Ulrich, Viktor Zouboulis, Paulina Sprezyna, Jasmin Schreiber, Tomer Landsberger, Maren Büttner, Moshe Biton, Eduardo J. Villablanca, Samuel Huber, Lorenz Adlung

## Abstract

Single-cell mRNA sequencing (scRNA-seq) allows deep molecular and cellular profiling of immunological processes. Longitudinal scRNA-seq datasets can be used for deterministic ordinary differential equation (ODE)-based modelling to mechanistically describe immune dynamics. Here, we derived longitudinal changes in the abundance of six colonic cell types during inflammatory bowel disease (IBD) from scRNA-seq data of a mouse model of colitis using ODE-based models. We then predicted the immune dynamics of a different mouse colitis protocol and confirmed these scRNA-seq-based predictions with our previously published single-cell-based flow cytometry data. We further hypothesised that the estimated model parameters reflect biological processes. We validated this prediction of cellular turnover rates with KI-67 staining and with gene expression information from the scRNA-seq data not used for model fitting. Finally, we tested the translational relevance of the model simulations by predicting genes indicative of treatment response in human IBD patients. The predictive power of IBD deterministic modelling from scRNA-seq data highlights its potential to advance our understanding of immune dynamics in health and disease.

## INTRODUCTION

The advent of single-cell mRNA-sequencing (scRNA-seq) technologies has elucidated various facets of the immune system, ultimately facilitating our understanding of disease aetiology as a dynamic process^1^. Despite the increasing availability of high-throughput data, its implementation in predictive dynamical models is hampered by insufficient temporal resolution. As a single measurement can capture multiple cellular states along a temporal continuum, many computational approaches seek to exploit this to reconstruct developmental trajectories^2^. Fate mapping and barcoding provide additional information to trace immune cellular histories^3^. Where omics datasets do have temporal resolution, statistical modelling frameworks are available to explore and analyse them^4–6^. However, when looking for associations between measured traits and experimental conditions, such statistical models treat time as a confounding factor.

In general, scRNA-seq technologies are suitable for the identification and in-depth molecular characterization of immune cell types and cellular states^7,8^. Transcriptional profiles reveal cellular heterogeneity, reflecting distinct functional states with potential clinical relevance^9^. The kinetics of these cell states can be used to dynamically model disease progression. While statistical models typically use data to infer disease trends and associations, dynamic models use deterministic equations to explicitly represent the underlying biological processes and predict disease dynamics based on established principles. Mathematical models in these cases describe changes in the abundance of cell populations over time based on a set of coupled ordinary differential equations (ODEs). To describe experimental data, parameters must be estimated that represent rate constants of biological processes such as proliferation, differentiation or death. ODE models are useful for simulating the dynamics of disease and discovering the principles governing in the progression of pathologies^10^.

Here, we used inflammatory bowel disease (IBD) as a paradigm for a dynamic immunological disease. IBD is characterised by relapsing-remitting inflammation of the gastrointestinal tract, associated with cramping, diarrhoea and weight loss. The two most common forms of IBD in humans are Crohn’s disease^11^ and ulcerative colitis^12^. Mouse models of the latter include T-cell transfer colitis and dextran sodium sulphate (DSS) colitis^13^. Omics technologies have great potential to inform the treatment of IBD patients through personalized medicine^14,15^. In this work, we explored scRNA-seq data from colitis in mice for deterministic modelling of IBD cell population dynamics. We aimed to create a generalizable ODE-based model indicative of biological processes during IBD and treatment response. We propose that such a systems biology approach can contribute to a better mechanistic understanding of immune-mediated inflammatory diseases.

## RESULTS

### Longitudinal single-cell mRNA-sequencing data reveals population dynamics during murine colitis

We obtained publicly available longitudinal scRNA-seq data from Ho *et al*. (GSE148794)^16^. The dataset contains 14,634 single-cell transcriptomes from the colon of mice (**Supplementary Fig. 1A**), which received 1.5% DSS in the drinking water for six days and afterwards water for another nine days. The DSS/water treatment regime represents a murine model of acute colitis followed by resolution of inflammation and mucosal healing in the colon of mice. Importantly, data was collected longitudinally during the experiment, meaning that mice were sacrificed and colon samples were collected at day 0, 3, 6, 9, 12, and 15. Per timepoint, three biological replicates (mice) were pooled and subjected to scRNA-seq by Smart-Seq2^17^. The mouse information was not hashed, but ten technical replicates were sequenced per time point, allowing us to estimate the uncertainty in the dynamics of cell population abundance (**Supplementary Fig. 1B**). We took cell types, as identified by Ho *et al*.^16^, and performed convolution to pseudo-bulk populations. This transformation allowed us to assess changes in colonic cell population abundance over time during DSS colitis. As a proof of concept, we selected six different cell populations for deterministic modelling. We chose “Abs. & sec cell”, which are epithelial cells (Epi); “B cell”, which are only B cells and no plasma cells; “Granulocyte”, which are described by Ho *et al*. as neutrophils (Neutr), because they all express S100a8/9^16^. In addition, we chose “MNPs”, which are mononuclear phagocytes, *i.e*., monocytes and macrophages (Mac); “Stromal”, which are stromal cells; and “T cell”, which are all T cells. Cell population annotations are summarized in **Table 1**.

**Table 1:** Cell population annotations.

| Annotation by<br>Ho *et al.*<sup>16</sup> | Description | Annotation<br>in this work |
| --- | --- | --- |
| Abs. & sec cell | Epithelial cells | Epi |
| B cell | B cells but not plasma cells | Bcell |
| Granulocyte | Granulocytes expressing S100a8/9, therefore Neutrophils | Neutr |
| MNPs | Mononuclear phagocytes: Monocytes + macrophages | Mac |
| Stromal | Stromal cells but not myofibroblasts | Stromal |
| T cell | All T cell subsets | Tcell |

We then used the extracted experimental data to fit an ODE model describing the change in abundance of each cell population over time during DSS colitis. Dynamic changes depend on parameters representing rate constants *k*. First, we tested different model structures by estimating rate constants given the experimental data. We then performed model reduction based on profile likelihood estimation^18^. Model selection was based on the corrected Akaike Information Criterion^19^ (**Supplementary Fig. 1C**). The best model was selected and included cell population specific turnover rates *k_turnover_*, as a ratio of proliferation to death (**Fig. 1A**). For example, the turnover rate *kB* of B cells is proportional to the accumulation of B cells in the mucosa and indirectly proportional to the depletion of B cells from the colonic tissue. In addition, the best model included a parameter representing mucosal healing, *kheal*. The rates of the best selected ODE model represent first order mass action kinetics and are shown in **Table 2**.

**Figure 1:**
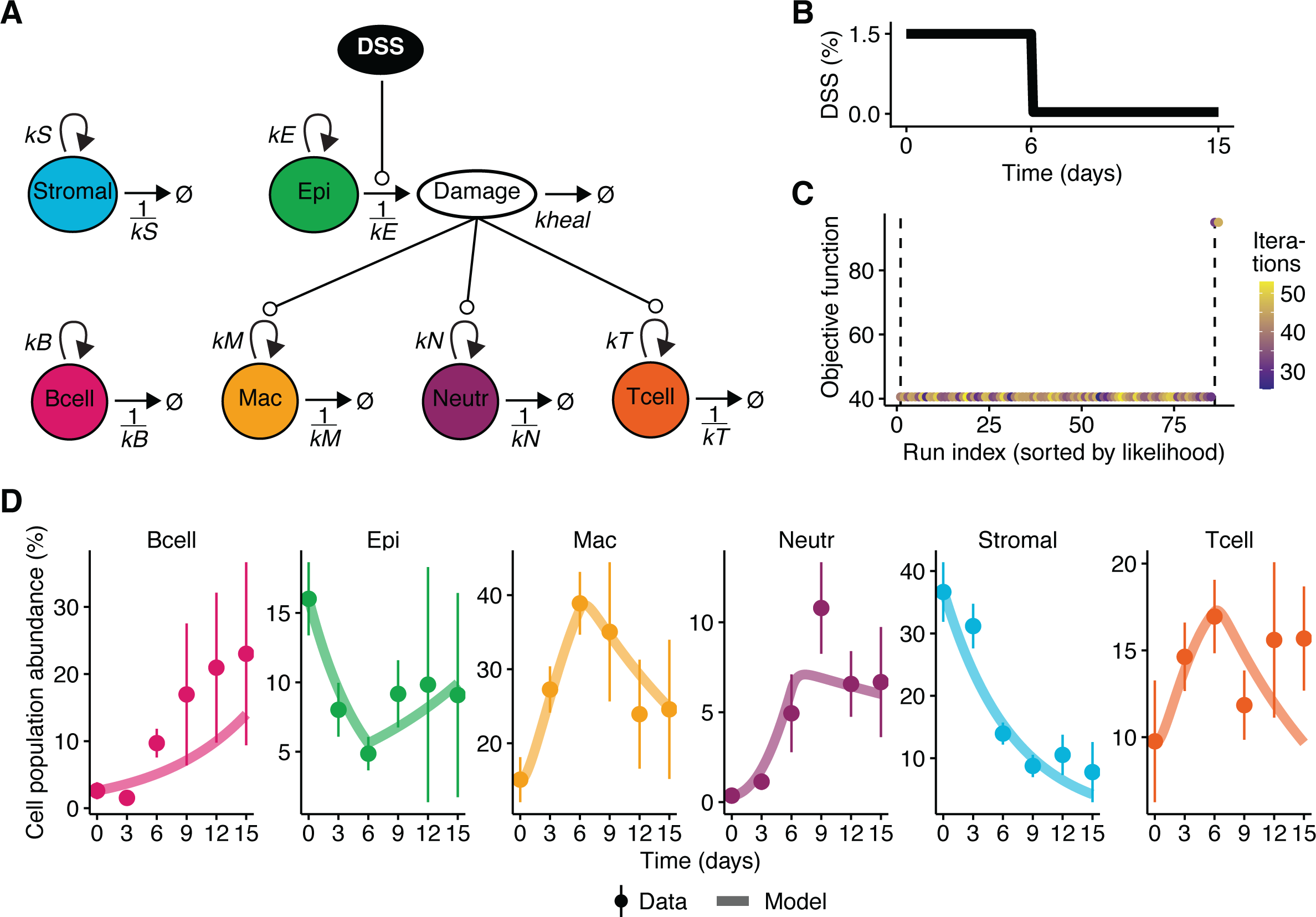
Model simulations, optimization of parameter estimation and model fit. **A** Schematic representation of the mathematical model. **B** DSS treatment as simulated model input. **C** Multi-start parameter estimation runs with likelihood as objective function. **D** Experimental data and simulations of the fitted model. Points represent mean values; error bars indicate standard deviations from experimental data. Solid lines represent model simulations.

**Table 2:** Rates of the best, selected ODE model.

|  | Source | -> | Sink | Rate | Description |
| --- | --- | --- | --- | --- | --- |
| 1 | | -> | Bcell | $k_B \cdot \text{Bcell}$ | B_Proliferation |
| 2 | Bcell | -> | | $(1/k_B) \cdot \text{Bcell}$ | B_Death |
| 3 | | -> | Mac | $\text{Damage} \cdot k_M \cdot \text{Mac}$ | Mac_Proliferaton |
| 4 | Mac | -> | | $(1/k_M) \cdot \text{Mac}$ | Mac_Death |
| 5 | | -> | Neutr | $\text{Damage} \cdot k_N \cdot \text{Neutr}$ | N_Proliferation |
| 6 | Neutr | -> | | $(1/k_N) \cdot \text{Neutr}$ | Neutr_Death |
| 7 | | -> | Tcell | $\text{Damage} \cdot k_T \cdot \text{Tcell}$ | T_Proliferation |
| 8 | Tcell | -> | | $(1/k_T) \cdot \text{Tcell}$ | T_Death |
| 9 | Stromal | -> | | $(1/k_S) \cdot \text{Stromal}$ | Stromal_Death |
| 10 | Epi | -> | Damage | $\text{DSS} \cdot (1/k_E) \cdot \text{Epi}$ | Epi_Death |
| 11 | | -> | Epi | $k_E \cdot \text{Epi}$ | Epi_Proliferation |
| 12 | Damage | -> | | $k_{heal} \cdot \text{Damage}$ | Healing |
| 13 | | -> | Stromal | $k_S \cdot \text{Stromal}$ | Stromal_Proliferation |
| 14 |  | -> | DSS | 0 | DSS |

DSS is supplied as input to the model. It is present at the given concentration for six days and absent for the remaining nine days of the experiment. The best model included DSS-induced epithelial breakdown and subsequent accumulation of ‘Damage’, which in turn attracts macrophages, T cells and neutrophils. Damage is resolved by mucosal healing. Our deterministic model proved to be fully structurally and practically identifiable^20^ (**Supplementary Fig. 1D**).

We then implemented the DSS treatment regimen (**Fig. 1B**) and used data from day 0 as initial values for model simulations and parameter estimation. A global optimum was found in 86% of the multi-start parameter estimation runs (**Fig. 1C**), indicating robust optimisation^21^. Our fitted model was able to capture the dynamics of cell population abundance extracted from the scRNA-seq dataset (**Fig. 1D**). B cells accumulated over time, whereas stromal cells declined. Macrophages, neutrophils, and T cells were initially recruited to the tissue and then their abundance decreased during mucosal healing, which was reflected in the recovery of epithelial cell abundance after the initial DSS-induced epithelial decay.

In summary, our selected deterministic model captures cell population abundance dynamics during murine colitis.

### Independent validation of the fitted ODE model with single cell data

We then sought to validate our fitted model with an independent single-cell-based, longitudinal dataset of murine DSS colitis: flow cytometry of colonic immune cells, which we have previously published^22^. We transformed cell counts per timepoint into relative frequencies and used information at day 0 within three standard deviations as initial values. We then predicted seven days of 2.5% DSS treatment followed by seven days of water using the parameters from our model fitted to the other dataset (*c.f*., **Fig. 1D**). There was a qualitatively good agreement between the model prediction and the experimental validation (**Fig. 2A**). It should be noted that the validation experiment is a completely independent study, even obtained by a different technology; scRNA-seq vs. flow cytometry. The experimental protocol, *i.e*., DSS concentration and timing, was different, which we simulated as input accordingly. In general, there are several statistical models to infer robust changes in cell type abundance from scRNA-seq data, *e.g*., scCODA^23^, or sccomp^24^. A continuous model for testing for differential abundance in scRNA-seq data is Milo^25^. Of note, none of these models are designed for time series data.

**Figure 2:**
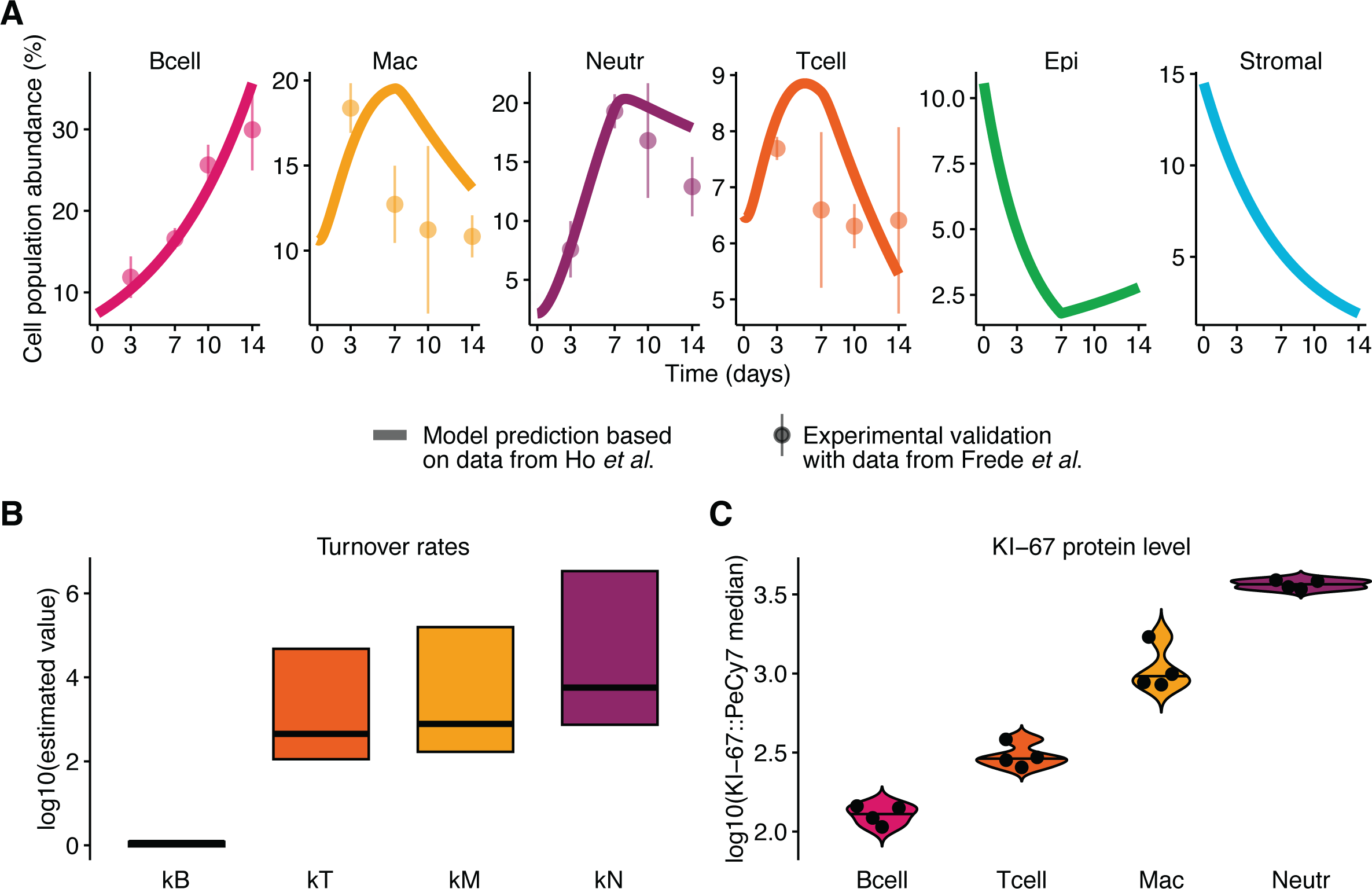
Independent validations of model predictions. **A** ODE model was fitted to single-cell mRNA-sequencing data from Ho *et al*.^16^ Estimated parameters of best model were chosen for model prediction. DSS input was adapted accordingly. Experimental validation by flow cytometry data from Frede *et al*.^22^ Initial values were varied within three standard deviations of experimental data. Lines represent model predictions. Points represent mean values, and error bars indicate standard deviation between each three mice of validation experiment. **B** Model prediction: 95% confidence bounds of estimated model parameters of best fit. Horizontal line within the confidence bounds indicates global optimum. **C** Violin plots of median KI-67 protein level of immune cell populations measured by flow cytometry. Every dot represents a mouse.

Overall, our fitted ODE-based model generalizes well to an independent dataset (**Supplementary Fig. 2A**, predicted vs. observed, Pearson’s rho=0.92). The good qualitative agreement between model prediction and experimental validation highlights the potential of the ODE model to predict immune dynamics when the model structure, initial abundance and its variance are known.

### Predictive power of an ODE model fitted to longitudinal scRNA-seq data

We then asked whether the estimated rate constants of the fitted ODE model were indicative of cellular gene expression. If this is the case, transcriptional programmes inferred from scRNA-seq data can potentially inform parameter estimation, *e.g*. by determining priors based on expression levels. For example, if the expression of a cell cycle score of a cell population is rather high in the scRNA-seq data, one would expect to find a high estimate of the rate constant for the proliferation parameter of the cell population in the ODE model. To test this hypothesis, we examined the confidence bounds of the optimised turnover rates per cell population (**Supplementary Fig. 2B**). These estimated parameters were used as model predictions. High turnover rates reflect the expansion of cell populations in the mucosa or *lamina propria* during DSS colitis (as change in percentage of cell type abundance per day). Immune cells with high turnover rates are therefore expected to be more pro-inflammatory, while non-immune cells with high turnover rates would imply mucosal healing under these conditions. We reasoned that a gene expression signature reflecting both intestinal inflammation and tissue damage would best correlate with the model prediction of estimated turnover rates. We selected the gene module of the KEGG pathway hsa05321: ‘Inflammatory bowel disease – Homo sapiens (human)’ for experimental validation. After converting the genes to their murine homologues, we calculated the IBD expression signature from the scRNA-seq data as experimental validation (**Supplementary Fig. 2C**). Note that gene expression was not used for model fitting. The estimated turnover rates and the calculated IBD expression score were found to be correlated (predicted vs. observed, Pearson’s rho=0.81). Model predictions of turnover rates were thus experimentally validated by expression of the IBD gene module.

As the IBD gene module contains 65 genes, we wondered whether a simpler physiological molecular readout would also reflect the estimated turnover rates. To test this hypothesis, we performed a dedicated experiment. We treated mice with 1.8% DSS as effective concentration for seven days and allowed them to recover with water for a further seven days. We then isolated colonic immune cells and stained them for the antigen KI-67 (**Supplementary Fig. 2D**), which serves as a proliferation marker^26^. Our flow cytometry data show that the estimated turnover rates of the immune cell populations (**Fig. 2B**) are indicative of KI-67 protein levels (**Fig. 2C**, **Supplementary Fig. 2E**, predicted vs. observed, Pearson’s rho=0.88). Taken together, our results indicate that estimated turnover rates can be informed by gene expression data and at the same time be predictive of biological processes.

### Translational implications of a fitted ODE model of murine colitis

Finally, we investigated whether the fitted ODE model reflected processes in human IBD. To this end, we reasoned that genes correlated with epithelial cell dynamics might be indicative of mucosal healing upon DSS retrieval. We therefore correlated epithelial cell abundance with pseudo-bulk gene expression per time point from Ho *et al*.^16^ Simulated epithelial cell abundance yields a different distribution of Pearson correlation coefficients than experimentally measured epithelial cell abundance, with more highly correlated genes on the right side of the distribution (**Supplementary Fig. 3A**). The number of significantly correlated genes (p<0.05) increased from 2,590 for measured epithelial abundance to 4,301 for simulated epithelial abundance. These results highlight the advantage of our fitted ODE model for simulating epithelial abundance, robustly linking it to pseudo-bulk gene expression. Among the most highly correlated genes were many genes associated with cell-cell contact, such as *Cldn8*, *Cftr*, *Crb3* and *Jam2*, as well as genes associated with ion exchange, such as *Slc26a3* and *Trpm6* (**Fig. 3A**). To test whether these genes are relevant to human IBD, we obtained publicly available Affymetrix microarray data from Arijs *et al*. (GSE16879)^ref.27^ from 61 IBD patients before and four to six weeks after their first infliximab infusion. We looked for differential gene expression in pre-treatment colon between responders and non-responders. We found genes such as *APQ8*, which encodes aquaporin 8, but also *CLDN8*, *TRPM6* and *SLC26A3* (**Fig. 3B**). To test the overlap more systematically between genes positively correlated with epithelial abundance in our mouse model and genes upregulated in IBD patients responding to infliximab treatment, we performed a hypergeometric test on the intersection between the two gene sets and obtained a significant overlap of 25 genes (**Fig. 3C**). These 25 genes were associated with KEGG pathways of physiological gut functions such as ‘tight junction’, ‘mineral absorption’ and ‘synthesis and degradation of keton bodies’ (**Fig. 3D**), suggesting that residual barrier function is an indicator of response to infliximab treatment.

**Figure 3:**
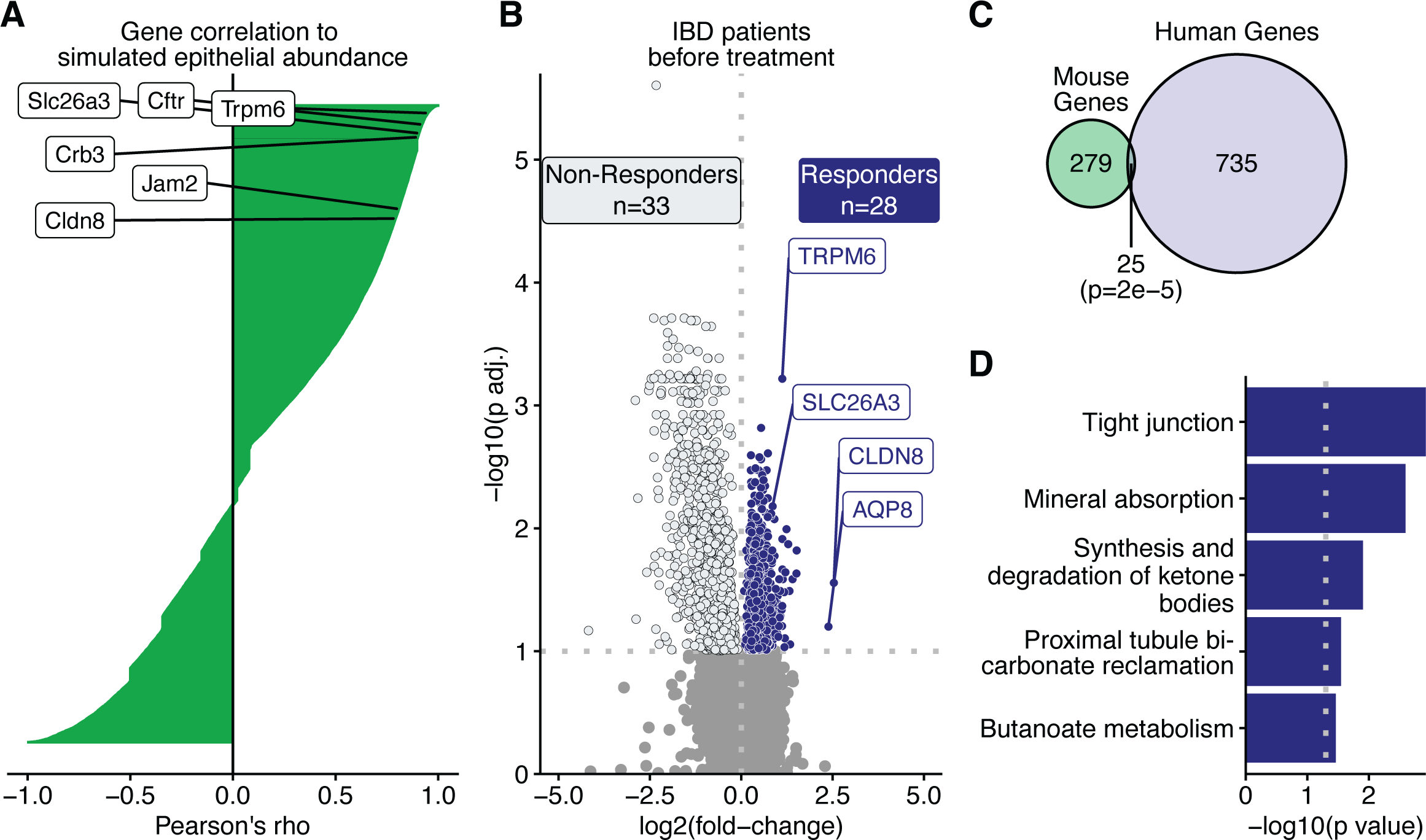
Testing of model-based gene predictions in human IBD. **A** Waterfall plot of genes ordered by their correlation to simulated epithelial abundance. Correlation coefficient is Pearson’s rho. Exemplary genes with high, positive correlation are given. **B** Vulcano plot of differentially expressed genes in colonic biopsies from IBD patient before treatment. Responders vs. Non-Responders. Exemplary genes are given. Mann-Whitney U test. p values adjusted for multiple testing. **C** Venn diagram of gene set overlap between mouse genes significantly positively correlating with simulated epithelial abundance and genes overexpressed in human IBD patients responding to treatment. Hypergeometric test. **D** Bar chart of KEGG pathways associated with overlapping genes from **C**.

Looking at genes negatively correlated with simulated epithelial abundance, we found pro-inflammatory genes such as *Il1b* and anti-inflammatory genes such as *Il4ra* and *Il10ra* among the most negatively correlated genes (**Supplementary Fig. 3B**). Conversely, *IL1B* and *IL10*, as well as *S100A9*, were upregulated in pre-treatment biopsies from non-responders (**Supplementary Fig. 3C**). The overlap between these two gene sets contained 73 genes (**Supplementary Fig. 3D**). These 73 genes were associated with KEGG pathways of inflammation such as ‘cytokine cytokine receptor interaction’ and ‘JAK STAT signaling pathway’ (**Supplementary Fig. 3E**), which warrants further investigation.

In conclusion, our fitted ODE model allows the simulation of epithelial abundance to identify genes that are indicative of mucosal healing and barrier integrity in mice, which may ultimately be associated with a positive response to infliximab treatment in human IBD patients or provide treatment alternatives.

## DISCUSSION

In this study, we explored the extent to which longitudinal scRNA-seq data can inform deterministic modelling of murine colitis. The amount of publicly available omics datasets is steadily increasing, and we expect more longitudinal studies to be published in the future. Here, we provide a first proof-of-concept study showing that Smart-Seq2 data can be used to infer cell population abundance dynamics. Although there is still considerable technical variability and information on biological variability are lacking (**Supplementary Fig. 1B**), the extracted information is sufficient to identify a set of seven kinetic parameters of an ODE model describing the change in colonic cell population abundance during murine colitis (**Supplementary Fig. 1D**). We originally developed ‘Data2Dynamics’^28^, an open source framework for deterministic modelling, but it only runs under the proprietary software MATLAB. Meanwhile, free and open R packages ‘dMod’ and ‘cODE’ are available with similar functionality. With this setup we were able to capture the extracted dynamics from Ho *et al*. (**Fig. 1D**). We validated a prediction of our scRNA-seq fitted ODE model with an independent flow cytometry measurement of murine colitis (**Fig. 2A**). As flow cytometry allows longitudinal single-cell-based measurements, we have recently used such data for deterministic modelling of erythroid fate decisions^29^. We further validated the estimated turnover rates of our fitted ODE model by comparing them with single-cell gene expression levels of an IBD gene module (**Supplementary Fig. 2B,C**) and KI-67 staining (**Fig. 2B,C**). Future research will show whether differentiation markers or other reporters are also suitable to provide information on turnover rates in murine colitis models that not only address net proliferation but also apoptosis^3,30^.

Studies on human IBD increasingly focus on transcriptomic readouts^31–35^. One of the major challenges will be the implementation of these data into mathematical models for robust inference of biological properties. We have previously shown that multiple colon biopsies from the same site of an individual ulcerative colitis patient yield similar cell type frequencies when subjected to scRNA-seq^36^. However, high inter-individual variability, cell type robustness, isolation protocols and technologies are major sources of measurement error. The former could be addressed with more sophisticated statistical models such as mixed-effect models of cell type specific pseudo bulks^37^. Another limitation in inferring cell abundance dynamics from scRNA-seq data with clinical relevance is the lack of spatial resolution. We anticipate that the sensitivity of the technologies will improve massively in the future, so that partial differential equation models will soon be able to infer cellular and molecular dynamics across space and time for systems immunology in IBD and beyond.

## METHODS

### Mice

All mice were of C57/BL6J wild-type background and bred in the UKE animal facility. Animals were kept in accordance with the institutional review board ‘Behörde für Soziales, Familie, Gesundheit und Verbraucherschutz’ (Hamburg, Germany), and under specific-pathogen-free conditions at ambient temperature. Experimental procedures were approved by the local ethics committee under the number N033/23. Mice were all male and between 8 and 12 weeks of age.

### DSS-induced murine colitis model

Murine colitis was induced by administration of 1.8% DSS (Alfa Aesar) as the effective concentration dissolved in drinking water *ad libitum* for seven days, followed by seven days of pure water. Mouse physiology and body weight were monitored regularly. After 14 days, the mice were sacrificed, and the colons were collected for further processing.

### Isolation of colonic cells

Excised colon samples were extensively washed of luminal content and opened longitudinally. The tissue was cut into 1 cm pieces and then incubated in HBSS (Gibco) containing 5 mM EDTA, 5% FBS, 10 mM HEPES and 1 mM DTT at 37°C with shaking for 20 minutes. Samples were then digested in HBSS with CaMg, 1mg/ml collagenase VIII (Sigma), 0.4mg/ml DNAase I, 5% FBS and 10mM HEPES at 37°C using the gentleMACS Octo Dissociator with Heaters’ programme ‘37C_m_LPDK_1’. Isolated cells were washed once in PBS (2mM EDTA, 1% FBS) and lymphocytes were enriched on a 40-67% Percoll gradient (GE Healthcare).

### Flow cytometry

Cells were stained with live/dead dye and CD45 (clone: 30-F11), CD90.2 (clone: 53-2.1), CD3 (clone: 17A2), B220 (clone: RA3-6B2), I-A/I-E (clone: M5/114.15.2), CD11b (clone: M1/70), CD64 (clone: X54-5/7.1) and, Ly-6G (clone: 1A8) surface antibodies at 4°C for 20 minutes.

After washing twice, intracellular staining for KI67 (clone: B56) was performed on fixed cells using the Foxp3/Transcription Factor Staining Buffer Set (eBioscience) according to the manufacturer’s instructions. Multiparameter analysis was performed on an LSR Fortessa II (BD) and analysed with FlowJo software (FlowJo, LLC).

### mRNA-sequencing analysis

scRNA-seq data was processed with the R package ‘Seurat’ (v. 4.3.0), convolution to pseudo-bulk was performed with its ‘AverageExpression’ function. Affymetrix microarray analysis was performed with the R packages ‘Biobase (v. 2.58.0) and ‘GEOquery’ (v. 2.66.0). Differential gene expression analysis of Affymetrix microarray data was performed by Mann-Whitney U test with false-discovery rate (FDR) correction. KEGG pathway and IDG drug target analysis was performed with the R package ‘enrichR’ (v. 3.2).

### Statistical analysis

Equality of variances in distributions of correlation coefficients was assessed by Levene’s test with the R package ‘car’ (v. 3.1-2). Hypergeometric test for overlap of two gene sets was performed with the R package ‘stats’ (v. 4.2.1).

### Deterministic modelling

Our deterministic model assumes that the change of abundance of cell populations 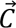 with time *t* during colitis can be described as a function of those cell populations 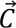 and estimated parameters 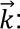

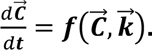

The ODE model was implemented with R packages ‘dMod’ and ‘cOde’, which are available at https://github.com/dkaschek/. Default settings were used for optimization runs, with *normL2* as objective function and priors with *mean*=1e0, *sigma*=1e1. 100 fits were run per optimization with *mstrust* optimizer. The objective function is the weighted sum of squared residuals:

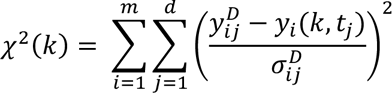

where 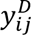 denotes *d* data points for all observables *i* at timepoint *t_j_*. 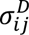 refers to the respective measurement error, and *y_i_*(*k*, *t_j_*) is the simulation of the observable *i* at timepoint *t_j_* as a function of the parameters *k*. Parameter estimation was performed by minimizing the objective function:

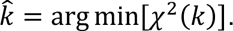

For normally distributed measurement error, this corresponds to the maximum likelihood estimate of *k*, in turn:

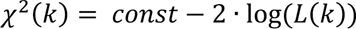

with *L*(*k*) represents the likelihood.

## Data and code availability

All data of this study and the code required to reproduce figures will be made freely available upon publication at: https://github.com/AdlungLab.

## ACKNOWLEDGEMENTS

All authors thank the AdlungLab members and Gemma Douilhet for fruitful discussions. C.K.: Supported by the iDfellows Hamburg Clinician Scientist Programme in Infectious Diseases (DFG, German Research Foundation, funding code: 493624519). L.A.: Supported by funding of the Klaus Tschira Boost Fund, a joint initiative of the German Scholars Organization and the Klaus Tschira Foundation, and the *Behörde für Wissenschaft, Forschung, Gleichstellung und Bezirke (BWFGB) der Freien und Hansestadt Hamburg* – Hamburg Innovation (HI) ‘Calls for Transfer’.

## AUTHOR CONTRIBUTIONS

C.K. performed computational modelling and revised the manuscript. H.U. performed validation experiments and revised the manuscript. V.Z. performed computational modelling and revised the manuscript. P.S. assisted with validation experiments. J.S., T.L., M.Bü and M.Bi. revised the manuscript. E.J.V. provided validation data and revised the manuscript. S.H. provided mice for validation experiments and revised the manuscript. L.A. performed computational modelling, conceived the study, analysed the data, drafted the manuscript, and revised the manuscript.

---

**Supplementary Figure 1:**
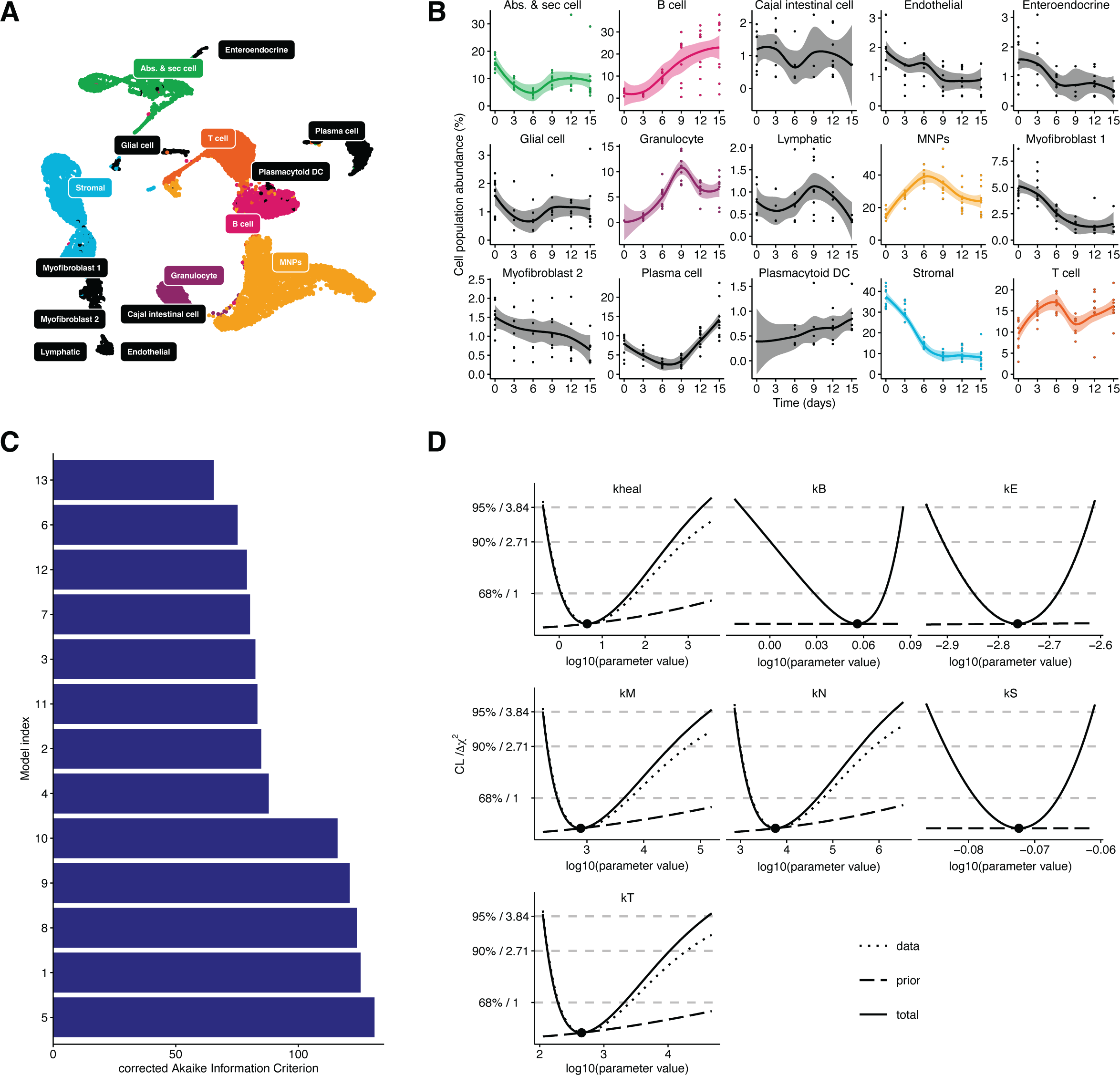
Data extraction and model fitting. **A** tSNE plot of single-cell mRNA-sequencing data (GSE148794) of 14,634 cells. **B** Extracted abundance of cell populations at different time points from ten technical replicates of each three pooled mice per time point. Colored populations were chosen for mathematical modelling. **C** Indices of different model structures fitted to experimental data and evaluated by the corrected Akaike Information Criterion. Model 13 was selected as best model. **D** Profile likelihood estimates of the parameters of the selected best model.

**Supplementary Figure 2:**
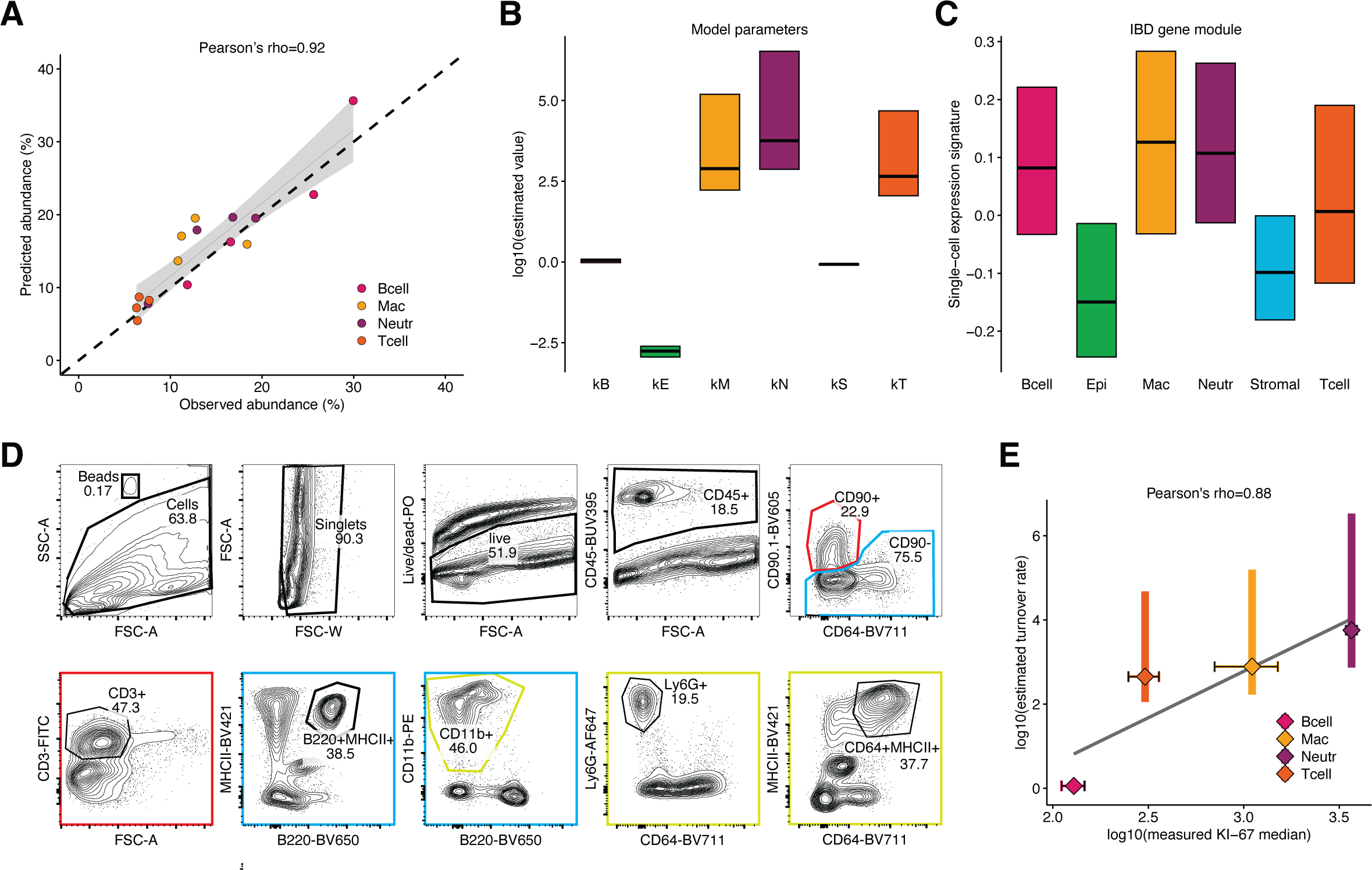
Experimental validations of model predictions. **A** Predicted cell population abundance based on data by Ho *et al*. vs. observed cell population abundance measured by Frede *et al*. Pearson’s rho is given. **B** Model prediction: 95% confidence bounds of estimated model parameters of best fit. Horizontal line within the confidence bounds indicates global minimum. **C** Experimental validation: Box plot of normalized gene expression score of IBD gene module (hsa05321) per single cell. Total of 12,500 cells. Upper line indicates 75% percentile, middle line indicates median, lower line indicates 25% percentile. **D** Gating scheme of the DSS validation experiment. **E** Estimated turnover rates based on data by Ho *et al*. vs. measured KI-67 median levels per colonic immune populations from DSS treated mice. Vertical bars indicate 95% confidence region. Horizontal error bars indicate standard deviation from each four mice. Solid grey line indicates linear fit. Pearson’s rho is given.

**Supplementary Figure 3:**
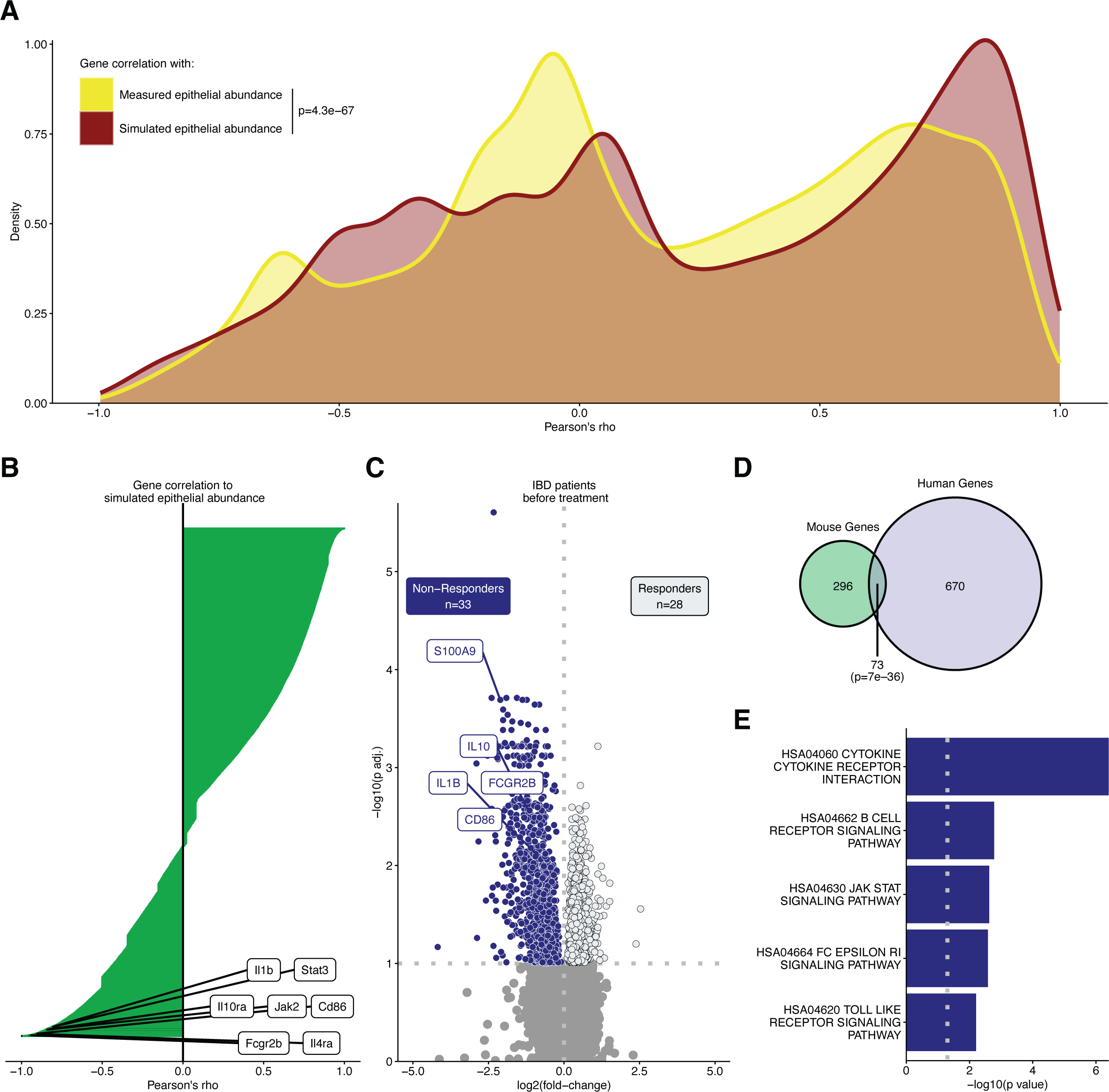
Comparison of measured and simulated epithelial abundance. **A** Distributions of gene-wise correlation to measured or simulated epithelial abundance. Correlation coefficient is Pearson’s rho. Levene’s test for equality of variance. **B** Waterfall plot of genes ordered by their correlation to simulated epithelial abundance. Correlation coefficient is Pearson’s rho. Exemplary genes with strong, negative correlation are given. **C** Vulcano plot of differentially expressed genes in colonic biopsies from IBD patient before treatment. Responders vs. Non-Responders. Exemplary genes are given. Mann-Whitney U test. p values adjusted for multiple testing. **D** Venn diagram of gene set overlap between mouse genes significantly negatively correlating with simulated epithelial abundance and genes overexpressed in human IBD patients not responding to treatment. Hypergeometric test. **E** Bar chart of KEGG pathways associated with overlapping genes from **D**.

## REFERENCES

1. Adlung, L. & Amit, I. From the Human Cell Atlas to dynamic immune maps in human disease. Nat. Rev. Immunol. 18, 597–598 (2018).

2. Ding, J., Sharon, N. & Bar-Joseph, Z. Temporal modelling using single-cell transcriptomics. Nat. Rev. Genet. 23, 355–368 (2022).

3. Höfer, T. & Rodewald, H.-R. Differentiation-based model of hematopoietic stem cell functions and lineage pathways. Blood 132, 1106–1113 (2018).

4. Mor, U. et al. Dimensionality reduction of longitudinal ’omics data using modern tensor factorizations. PLOS Comput. Biol. 18, e1010212 (2022).

5. Velten, B. et al. Identifying temporal and spatial patterns of variation from multimodal data using MEFISTO. Nat. Methods 19, 179–186 (2022).

6. Vasaikar, S. V. et al. A comprehensive platform for analyzing longitudinal multi-omics data. Nat. Commun. 14, 1684 (2023).

7. Bonaguro, L. et al. A guide to systems-level immunomics. Nat. Immunol. 23, 1412–1423 (2022).

8. Velten, B. & Stegle, O. Principles and challenges of modeling temporal and spatial omics data. Nat. Methods 20, 1462–1474 (2023).

9. Papalexi, E. & Satija, R. Single-cell RNA sequencing to explore immune cell heterogeneity. Nat. Rev. Immunol. 18, 35–45 (2018).

10. Yue, R. & Dutta, A. Computational systems biology in disease modeling and control, review and perspectives. Npj Syst. Biol. Appl. 8, 1–16 (2022).

11. Roda, G. et al. Crohn’s disease. Nat. Rev. Dis. Primer 6, 22 (2020).

12. Kobayashi, T. et al. Ulcerative colitis. Nat. Rev. Dis. Primer 6, 74 (2020).

13. Katsandegwaza, B., Horsnell, W. & Smith, K. Inflammatory Bowel Disease: A Review of Pre-Clinical Murine Models of Human Disease. Int. J. Mol. Sci. 23, 9344 (2022).

14. Zheng, H. B. Application of single-cell omics in inflammatory bowel disease. World J. Gastroenterol. 29, 4397–4404 (2023).

15. Agrawal, M., Allin, K. H., Petralia, F., Colombel, J.-F. & Jess, T. Multiomics to elucidate inflammatory bowel disease risk factors and pathways. Nat. Rev. Gastroenterol. Hepatol. 19, 399–409 (2022).

16. Ho, Y.-T. et al. Longitudinal Single-Cell Transcriptomics Reveals a Role for Serpina3n-Mediated Resolution of Inflammation in a Mouse Colitis Model. Cell. Mol. Gastroenterol. Hepatol. 12, 547–566 (2021).

17. Picelli, S. et al. Smart-seq2 for sensitive full-length transcriptome profiling in single cells. Nat. Methods 10, 1096–1098 (2013).

18. Maiwald, T. et al. Driving the Model to Its Limit: Profile Likelihood Based Model Reduction. PLOS ONE 11, e0162366 (2016).

19. Model Selection and Multimodel Inference. (Springer New York, 2004). doi:10.1007/b97636.

20. Raue, A. et al. Structural and practical identifiability analysis of partially observed dynamical models by exploiting the profile likelihood. Bioinformatics 25, 1923–1929 (2009).

21. Raue, A. et al. Lessons Learned from Quantitative Dynamical Modeling in Systems Biology. PLOS ONE 8, e74335 (2013).

22. Frede, A. et al. B cell expansion hinders the stroma-epithelium regenerative cross talk during mucosal healing. Immunity 55, 2336–2351.e12 (2022).

23. Büttner, M., Ostner, J., Müller, C. L., Theis, F. J. & Schubert, B. scCODA is a Bayesian model for compositional single-cell data analysis. Nat. Commun. 12, 6876 (2021).

24. Mangiola, S. et al. sccomp: Robust differential composition and variability analysis for single-cell data. Proc. Natl. Acad. Sci. U. S. A. 120, e2203828120 (2023).

25. Dann, E., Henderson, N. C., Teichmann, S. A., Morgan, M. D. & Marioni, J. C. Milo: differential abundance testing on single-cell data using k-NN graphs. 2020.11.23.393769 Preprint at 10.1101/2020.11.23.393769 (2020).

26. Gerdes, J., Schwab, U., Lemke, H. & Stein, H. Production of a mouse monoclonal antibody reactive with a human nuclear antigen associated with cell proliferation. Int. J. Cancer 31, 13–20 (1983).

27. Arijs, I. et al. Mucosal Gene Expression of Antimicrobial Peptides in Inflammatory Bowel Disease Before and After First Infliximab Treatment. PLOS ONE 4, e7984 (2009).

28. Raue, A. et al. Data2Dynamics: a modeling environment tailored to parameter estimation in dynamical systems. Bioinforma. Oxf. Engl. 31, 3558–3560 (2015).

29. Adlung, L. et al. Cell-to-cell variability in JAK2/STAT5 pathway components and cytoplasmic volumes defines survival threshold in erythroid progenitor cells. Cell Rep. 36, (2021).

30. Patankar, J. V. & Becker, C. Cell death in the gut epithelium and implications for chronic inflammation. Nat. Rev. Gastroenterol. Hepatol. 17, 543–556 (2020).

31. Thomas, T. et al. A longitudinal single-cell therapeutic atlas of anti-tumour necrosis factor treatment in inflammatory bowel disease. 2023.05.05.539635 Preprint at 10.1101/2023.05.05.539635 (2023).

32. Millet, V. et al. Harnessing the Vnn1 pantetheinase pathway boosts short chain fatty acids production and mucosal protection in colitis. Gut 72, 1115–1128 (2023).

33. Maddipatla, S. C. et al. Assessing Cellular and Transcriptional Diversity of Ileal Mucosa Among Treatment-Naïve and Treated Crohn’s Disease. Inflamm. Bowel Dis. 29, 274–285 (2023).

34. Argmann, C. et al. Biopsy and blood-based molecular biomarker of inflammation in IBD. Gut 72, 1271–1287 (2023).

35. Martin, J. C. et al. Single-Cell Analysis of Crohn’s Disease Lesions Identifies a Pathogenic Cellular Module Associated with Resistance to Anti-TNF Therapy. Cell 178, 1493–1508.e20 (2019).

36. Smillie, C. S. et al. Intra- and Inter-cellular Rewiring of the Human Colon during Ulcerative Colitis. Cell 178, 714–730.e22 (2019).

37. Chen, M. & Dahl, A. A robust model for cell type-specific interindividual variation in single-cell RNA sequencing data. 2023.02.24.529987 Preprint at 10.1101/2023.02.24.529987 (2023).

